# Kinome expansion in the *Fusarium oxysporum* species complex driven by accessary chromosomes

**DOI:** 10.1101/308064

**Authors:** Gregory A. DeIulio, Li Guo, Yong Zhang, Jonathan Goldberg, H. Corby Kistler, Li-Jun Ma

## Abstract

The *Fusarium oxysporum* species complex (FOSC) is a group of soil-borne pathogens causing severe disease in over one hundred plant hosts, while individual strains exhibit strong host specificity. Both chromosome transfer and comparative genomics experiments have demonstrated that lineage-specific (LS) chromosomes contribute to the host specific pathogenicity. However, little is known about the functional importance of genes encoded in these LS chromosomes. Focusing on signaling transduction, this study compared kinomes of 12 *F. oxysporum* isolates, including both plant and human pathogens and one non-pathogenic biocontrol strain, with seven additional publicly available ascomycete genomes. Overall, *F. oxysporum* kinomes are the largest, facilitated in part by the acquisitions of the LS chromosomes. The comparative study identified 99 kinases that are present in almost all examined fungal genomes, forming the core signaling network of ascomycete fungi. Compared to the conserved ascomycete kinome, the expansion of the *F. oxysporum* kinome occurs in several kinases families such as Histidine kinases that are involved in environmental signal sensing and TOR kinase that mediates cellular responses. Comparative kinome analysis suggests a convergent evolution that shapes individual *F. oxysporum* isolates with an enhanced and unique capacity for environmental perception and associated downstream responses.

**IMPORTANCE:** Isolates of *F. oxysporum* are adapted to survive a wide range of host and non-host conditions. In addition, *F. oxysporum* was recently recognized as the top emerging opportunistic fungal pathogen infecting immunocompromised humans. The sensory and response networks of these fungi undoubtedly play a fundamental role in establishing the adaptability of this group. We have examined the kinomes of 12 *F. oxysporum* isolates and highlighted kinase families that distinguish *F. oxysporum* from other fungi, as well as different isolates from one another. The amplification of kinases involved in environmental signal relay and regulating downstream cellular responses clearly sets *Fusarium* apart from other Ascomycetes. Though the function of many of these kinases is still unclear, their specific proliferation highlights them as a result of the evolutionary forces which have shaped this species complex, and clearly marks them as targets for exploitation in order to combat disease.

## INTRODUCTION

A constantly evolving genome provides the genetic foundation for an organism to adapt to challenging environments. The species complex of *Fusarium oxysporum* represents an exceptional model to study the relationship between genome evolution and organism adaptation. Phylogenetically related, members within the *Fusarium oxysporum* species complex (FOSC) include both plant and human pathogens and are collectively capable of causing plant wilt diseases in over one hundred plant species. Individual isolates often exhibit a high degree of host specificity, reflecting rapid adaptation to particular host environments in a very short evolutionary time frame (in less than 30 Mya) (1). *Forma specialis* has been used to describe strains that are adapted to a specific host. Comparative studies revealed that horizontally acquired lineage-specific (LS) chromosomes contribute to the host-specific pathogenicity of each *forma specialis* (2–4).

Kinases are key regulators within cellular regulatory networks. They transduce extracellular and intracellular signals by modifying the activity of other proteins including transcription factors, enzymes, and other kinases via phosphorylation (5–7). A “protein kinome” encompasses all protein kinases within a genome (8–13) and can be divided into several families, including the STE (homologs of the yeast Sterile kinases), CK1 (Casein kinase 1), CAMK (Ca^2+^/calmodulin-dependent protein kinase), CMGC (cyclin-dependent kinases (<u>C</u>DKs), AGC (protein kinase A, G, and C families), HisK (Histidine kinase), Other, and Atypical families (14). Some fungal kinomes have been functionally characterized (15–21).

To study the variation and evolution of kinases with respect to FOSC host-specific adaptation, we compared kinomes of twelve *Fusarium oxysporum* isolates including ten plant pathogens, one human pathogenic strain, and one non-pathogenic biocontrol strain. In addition, we have included seven ascomycete fungal genomes available in public domains. Our study revealed a clear correlation between the genome size and the size of the kinome of an organism. Due in part to the acquisition of LS chromosomes, the sizes of FOSC genomes are larger than other genomes included in this study, and so are their kinomes. Regardless of kinome size, we observed a highly conserved kinome core of 99 kinases among all fungal genomes examined. In contrast to this remarkably stable core, variation among families and sub-families was observed across levels of taxonomic classification. Most interestingly, we observed the expansion of the target of rapamycin (TOR) kinase and histidine kinases among FOSC genomes. Monitoring nutrient availability and integrating intracellular and extracellular signals, the TOR kinase and its associated complex serves as a central regulator of cell cycle, growth, proliferation and survival (22). Increasing copy number of certain kinases may enable new functions, or add new temporal variations to existing pathways. The repeated, but independent, expansion of certain kinase families among FOSC genomes may suggest a fine tuning of similar pathways in responding to different host defenses or abiotic environmental challenges.

## RESULTS

### 1) Overall kinase conservation defines a core kinome among ascomycete fungi

We compared 19 Ascomycete fungal genomes (Table 1), including 12 strains within the FOSC, two sister species close to *F. oxysporum* (*F. graminearum* and *F. verticillioides*) (Figure 1a), two yeast genomes (*Saccharomyces cerevisiae* and *Schizosaccharomyces pombe*), two model fungal species (*Neurospora crassa* and *Aspergillus nidulans*), and an additional plant pathogen, *Magnaporthe oryzae*. With the exception of the two yeasts, other genomes were annotated at the Broad institute using the same genomic annotation pipeline (23). All kinomes were predicted using the kinome prediction pipeline (24) with the same parameters. Kinases were classified by Kinannote into the STE, CK1, CAMK, CMGC, AGC, HisK, Other, or Atypical families (Figure 1b, Supp. Table 1). Kinases that could not be classified within specified statistical parameters were categorized as unclassified.

**Figure 1.**
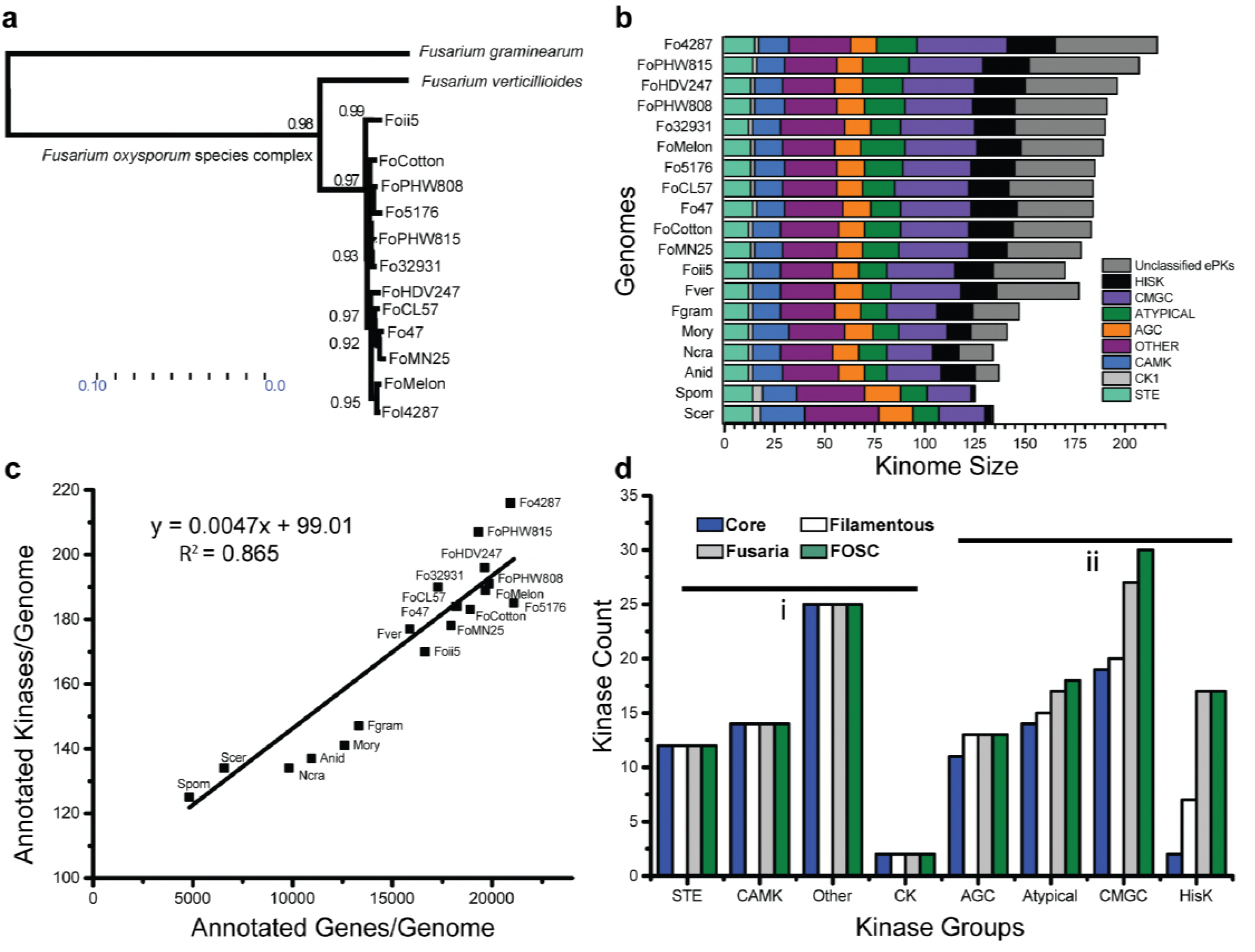
Kinomes Ascross Ascomycota. (a) A neighbor joining tree constructed from conserved genome genes showing the phylogenetic relationship among the Fusaria used in this study. (b) Kinases were broken up into major families and abundance per family plotted per species. Colors correspond to individual groups. (c) Total gene count plotted against kinome size in each genome. Origin coordinates are y=100, x=2500. (d) By eliminating kinases missing from more than one species, we compiled “conserved kinomes” for Ascomycetes (all fungi here), Filamentous (all but S. cerevisiae and S. pombe), the genus Fusaria, and the FOSC. Some families remain relatively stable across species (group i; AGC, CAMK, CK, Other, STE) while others expand due to increased copy number or increased sub-family number (group ii; Atypical, CMGC, HisK). Total size of conserved kinomes: Ascomycete (99), Filamentous (108), Fusaria (126), FOSC (128).

**Table 1.**
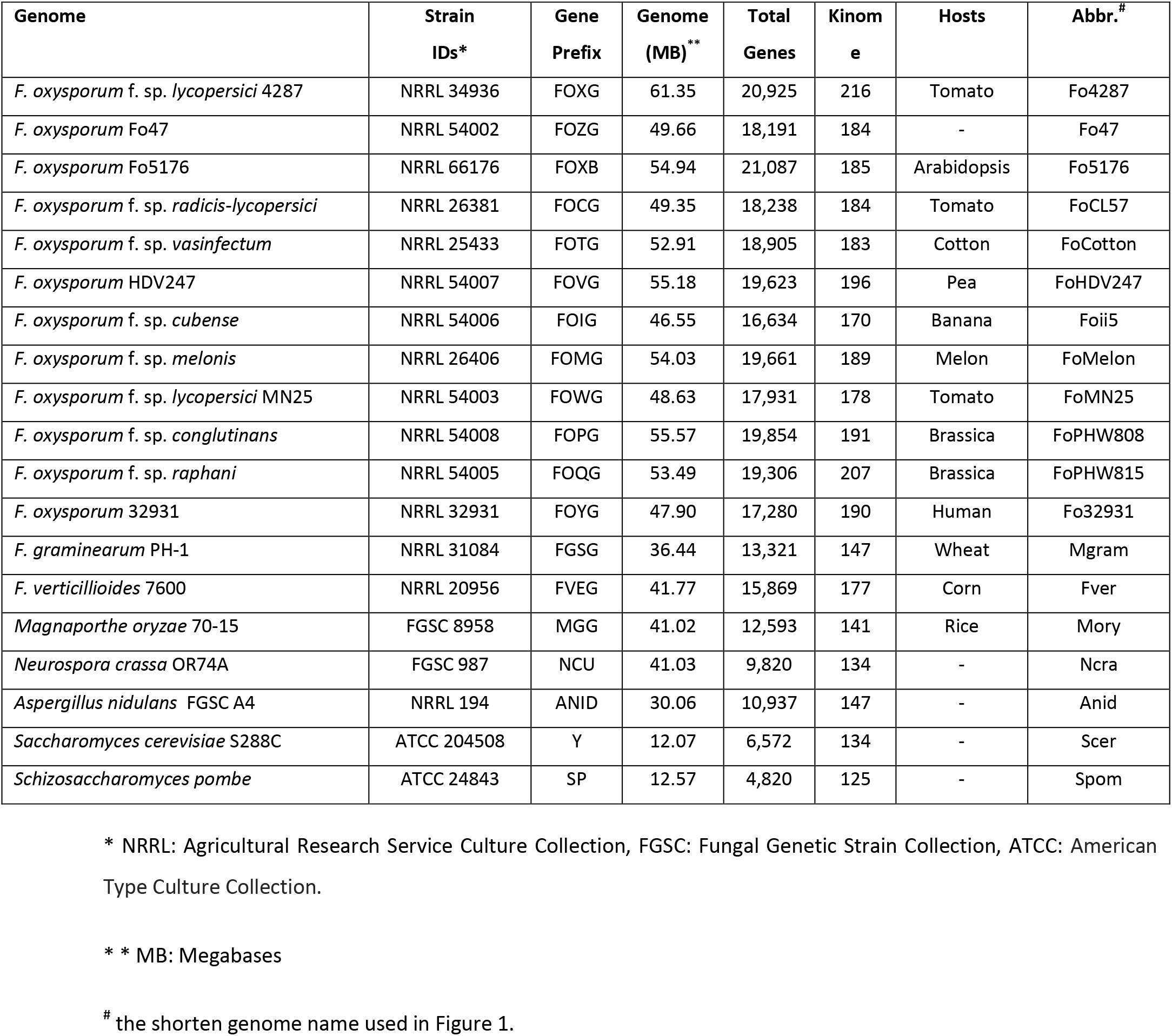
Fungal genomes used in this study.

Overall, there is a positive correlation between the total number of proteins encoded in a genome and the total number of protein kinases within that genome (y = 0.00473x + 99.01 R^2^= 0.86) (Figure 1c). The dependency of these two variables (number of kinases versus number of proteins) also points to a potential minimal number of 99 kinases for each ascomycete genome. Independently, we identified a core ascomycete kinome based on sequence conservation, taking a conservative approach by excluding kinase subfamilies that were missing from more than a single species. This conserved core kinome has all major kinase families, including 12 kinases in STE, 2 CK1, 14 CAMK, 11 AGC, 19 CMGC, 2 HisK, 14 Atypical, and 25 Other families (Figure 1d), totaling 99 conserved kinase orthologs, in agreement with the prediction of the regression model. In addition to the ascomycete core kinome, we also compiled conserved kinomes for filamentous fungi (excluding both *S. cerevisiae* and *S. pombe*), the genus Fusaria, and the FOSC, each containing 108, 127 and 131 kinases respectively. Interestingly, the sizes of the STE, CK1, CAMK, and Other families remain effectively constant across phylogenetic divisions, while AGC, Atypical, CGMC and HisK families exhibit trends of continuous expansion, illustrated as categories i and ii in Figure 1d.

#### The category I (constant group) includes STE, CK CAMK, and Other families

**STE family:** The STE kinases transduce diverse signals including osmolality, pheromone recognition and cell wall integrity and alter cell growth patterns in response to extracellular changes (25). The family includes MAP kinase kinases (MAPKK), MAP kinase kinase kinases (MAPKKK), and their upstream activators that function in MAPK signaling cascades. Evolutionarily conserved, a MAPKKK phosphorylates a MAPKK, and activated MAPKK phosphorylates a MAPK; activated MAPK phosphorylate other proteins to further control gene expression and cellular function. The 12 conserved fungal STE kinases are divided into nine subfamilies with a single kinase each, except the STE/PAKA and STE/YSK subfamilies that have two and three kinases respectively (Supp. Table 2). Except *S. cerevisiae*, which lacks a single STE/YSK ortholog, all other genomes contain 12 orthologs. Limited copy number variation of the STE family confirms the functional importance of these signaling pathways across Ascomycota.

**CK and CAMK families:** Like the STE kinases, the CK and CAMK kinases are highly conserved among the ascomycete fungal genomes. Each genome has 2 CK1 and 14 CAMK conserved subfamilies respectively. All filamentous fungal genomes have a single copy of each sub-family. The *S. pombe* genome lacks an ortholog of both the CAMK/CMK and the CAMK/Rad53 kinases, while the *S. cerevisiae* genome lacks an ortholog of the CAMK/CAMK1 kinase. An ancient kinase family of serine/threonine-selective enzymes, CK1 kinases are present in most eukaryotic organisms and are involved in important signal transduction pathways, including regulating DNA replication and the circadian-rhythm. In *S. cerevisiae*, the two CK1 kinases function in morphogenesis, proper septin assembly, endocytic trafficking, and glucose sensing (26)). Found in almost all eukaryotic cells, CAMK kinases are activated by Ca^2+^ fluctuations within the cell and control tip growth, branching, spore production, cell cycle progression, and secretion, among others (27).

**Other kinase family:** The Other kinase family includes several unique eukaryotic protein kinases (ePKs) that cannot be placed into any of the major ePK groups based on sequence similarity. There are 25 Other kinases, which are further divided into 24 subfamiles. Except for the Other/CAMKK subfamily, which contains two kinases, each subfamily of the Other kinase group has a single copy within each genome. The Other kinases account for a quarter of the ascomycete core kinome and many kinases within this group are involved in basal cellular functions including cell cycle control (28–30).

#### The category II (variable group) contains AGC, Atypical, CMGC, and HisKs

**AGC family:** The AGC family contains 11 subfamilies with a single orthologous copy each. All subfamilies are present in all genomes, except the AGC/YANK kinase is absent from the *S. cerevisiae* genome. AGC family kinases are cytoplasmic serine/threonine kinases regulated by secondary messengers such as cyclic AMP (PKA) or lipids (PKC) (31). They are involved in signaling pathways that orchestrate growth and morphogenesis, as well as response to nutrient limitation and other environmental stresses (32).

**Atypical kinase family:** There are 11 Atypical kinases, which are further divided into 10 subfamilies with a single copy in each subfamily, except the Atypical/ABC-1B subfamily which has two. Distinctively, kinases of the atypical family lack the canonical ePK domain, but have protein kinase activity (13). This family also includes many functionally important kinases including the TOR kinase, a major regulatory hub within the cell controlling nutrient sensing, cell cycle progression, stress responses, protein biosynthesis, and various mitochondrial functions (33). Other subfamilies of this group also play significant roles in cell cycle progression (RIO), mRNA degradation (PAN), and the DNA damage response (ATM, ATR, TRRAP).

**CMGC family:** The CMGC family is an essential and large group of kinases found in all eukaryotes, accounting for roughly 20% of most kinomes. The group is comprised of diverse subfamilies that control cell cycle, transcription, as well as kinases involved in splicing and metabolic control (34), including cyclin-dependent kinases (CDKs), mitogen-activated protein kinases (MAP kinases), glycogen synthase kinases (GSK), serine/arginine protein kinase (SRPK), and CDK-like kinases. Mitogen-activated protein kinases (MAPK) form the last step in the three step MAPK signaling cascades, which regulate functions from mating and invasive growth, to osmosensing and cell wall integrity (35). Cyclin-dependent kinases (CDK) are widely known as controllers of the cell cycle and transcription (34). Kinannote reported 23 subfamilies of CMGC kinase, of which 18 were conserved and included in the conserved kinome. Each subfamily contains a single kinase, except for the CMGC/ERK1 subfamily that contributes two. The *S. cerevisiae* kinome lacks 3 out of the 18 conserved subfamilies (CMGC/CDK11, CMGC/DYRK2, CMGC/PRP4).

**HisK family:** Differing from other kinase groups discussed above, the HisK family is widely distributed throughout prokaryotes and eukaryotes outside the metazoans. This group of kinases sense and transduce many intra– and extracellular signals (36–39). Distinctively, this family has the most significant expansion from yeast to filamentous fungi and is further expanded in the FOSC genomes (Figure 1d). All HisKs are classified into 11 families, or classes, based on the conservation of the H-box domain (36). However, only two classes (class V and class X) can be considered orthologous among most ascomycete genomes, present in all genomes here except *S. cerevisiae* (Supp. Table 2). The class V HisK is orthologous to the *S. pombe* Mak1, while the single conserved class X kinase is orthologous to the *S. pombe* Mak2/Mak3 kinases. Mak2 and Mak3 are known peroxide sensors (40), while all Mak1/2/3 are predicted to have a central role controlling the stress responses network (41).

### 2) LS chromosomes contribute to the individual expansion of FOSC kinomes

The conserved kinome represents defining pathways shared among Ascomycetes across evolutionary time. However, it is the unique additions to each kinome which enable species-specific adaptation. Among species here *F. oxysporum* kinomes are the largest. At roughly twice the size of the core ascomycete kinome, the average *Fusarium* kinome contains 185 kinases in 112 subfamilies (Table 1, Supp. Table 2). In addition, we observed a positive correlation between kinases encoded the LS region of each strain, defined as LS kinases here after, with the total number of LS genes (y = 0.0049x + 6.11, R^2^=0.57) (Supp. Fig 1a), suggesting that linage specific (LS) chromosomes contribute directly to the expanded kinomes. This correlation shares a similar slope as that of the whole genome, but effectively without a minimum kinase requirement.

In agreement with a comparative study among three *Fusarium* genomes (Ma, Does et al. 2010), little conservation was observed among the LS genomes (Supp. Figure 1b). The genome of each FOSC strain was partitioned into core and lineage specific (LS) using an “eliminating core” method (See Methods for detail). On average, the coverage of shared sequences among core genomes is over 70% (> 90% sequence similarity), as illustrated by supercontig 14 from the core. The average overlap for LS chromosomes of the reference strain 4287 was 6.3%. The most significant conservation was observed in chromosome 14 between two tomato pathogens MN25 (race 3) and the reference genome Fo4287 (race 2) (Supp. Table 3). This closeness was the result of the transfer of a pathogenicity chromosome, chromosome 14, suggested by comparative studies (2, 42). Other conserved fragments also observed, however their functional significance remains elusive.

To evaluate the overall functional importance of LS kinases, we have sequenced mRNA isolated from the reference genome Fo4287 under two experimental conditions, one at room temperature and the other shifted to 37 °C (Methods for details) (Supp. Table 5). Roughly half of the annotated Fo4287 genes (9,914) were expressed in either condition, including 85% of kinases within the conserved ascomycete kinome (85 out of the total 99 core kinases) (Supp. Table 6). In contrast, among the 44 LS kinases of Fo4287, the expression of only 8 (18%) was detected. Expressed LS kinases include 2 HisK (FOXG_14953, FOXG_15045), 2 Atypical/FunK1 (FOXG_12507, FOXG_14032), 2 Other/HAL (FOXG_06573, FOXG_07253), 1 STE/YSK (FOXG_14024), and 1 Unclassified kinase (FOXG_16175). For most cases, we saw higher levels of expression for the copies in the core compared to the LS copies.

Interestingly, heat stress at 37°C uniquely induced the expression of 4,394 genes, accounting for 44% of all expressed genes (Supp. Table 7). Similarly, about 44% of all expressed core kinases are induced under the heat stress, consistent with the functional conservation of the core genome. In contrast, 6 out of 8 expressed LS kinases were induced at 37°C, including a HAL kinase (FOXG_06573), a Class IV HisK (FOXG_14953), one Atypical/FunK1 (FOXG_12507) and three Unclassified kinases. The overall expression pattern of the Fo4287 kinome supports the potential function of some LS kinases, especially in coping with different stresses. Additional functional studies under diverse stress conditions may capture the expression of other, potentially condition specific, LS kinases.

### 3) Expanded FOSC kinases enhance signaling transduction in cell cycle control and environmental sensing

Expanded families belong to the category ii families (Figure 1d), enhancing functions related to cell cycle control and environmental sensing. For a soil-borne pathogen with strong host-specificity, like *F. oxysporum*, the adjustment of growth and cell cycle control in response to environmental cues are likely essential for survival.

#### 3.1 Enhanced cell cycle control centering on the target of rapamycin (TOR) kinase (Figure 2)

One of the most interesting expansions is the TOR kinase, which occurred in 7 out of 10 sequenced plant pathogenic FOSC strains. The TOR kinase is a top regulator of nutrient-sensing which dictates cellular responses according to the levels of nutrients and oxygen (43). A member of the Atypical/FRAP kinases, the TOR kinase is highly conserved in nearly all eukaryotic organisms from fungi to humans with few exceptions (44). Even though there are two TOR paralogs in the *S. cerevisiae* and the *S. pombe* genomes, almost all euascomycete fungal genomes have only one copy (Teichert et al., 2006). Our study revealed a TOR kinase expansion in 7 out of 12 sequenced FOSC strains, with 5 strains containing 1 and two strains containing 2 additional copies in addition to the single orthologous TOR kinase (Figure 2a). The orthologous copies of the TOR kinase (Marked as Core clade in Figure 2a) form a monophyletic group with almost identical amino acid sequences (>99.9%). A total of nine TOR paralogs (marked as LS clade in Figure 2a) clustered together with an average 95.9% amino acid identity compared to the orthologous copies, and an average 97% among LS paralogs (Supp. Figure 2), arguing against a recent duplication within an individual genome as a mechanism for the expansion of this gene family. However, there does appear to have been a recent duplication of one pair of the paralogs in a pea pathogenic strain (FOVG_18014 and FOVG_19124).

**Figure 2:**
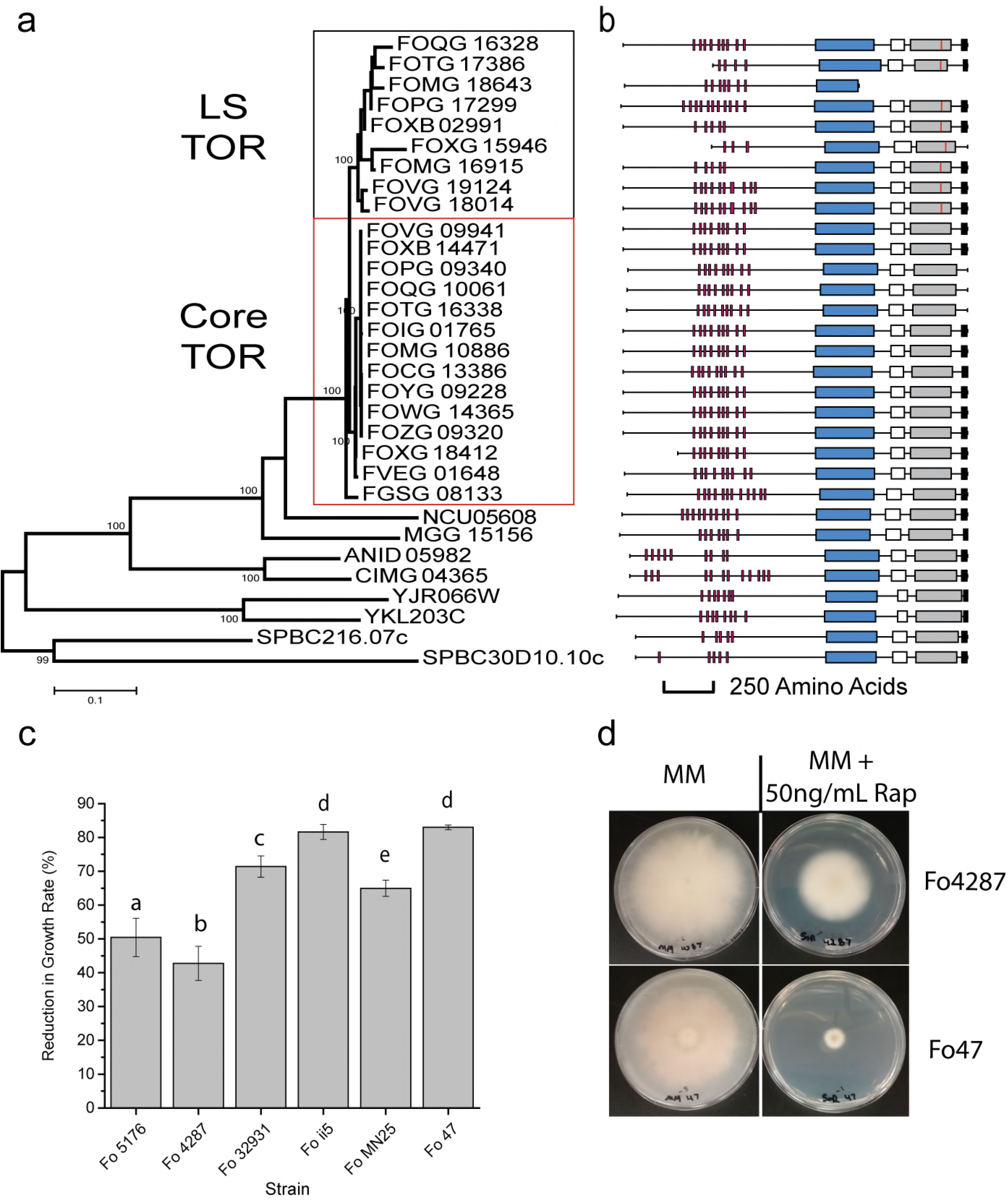
TOR kinase expansion through the LS genome. (a) A phylogenetic tree of TOR kinases constructed by NJ algorithm using TOR kinase protein sequence alignment based on the kinase domain. The core and LS clades of TOR kinase in Fo are shown in boxes. (b) On the right of the tree is a protein domain image showing the relative structure of each TOR kinase. A red vertical line inside the kinase domain indicates the mutation M2345L. Domain shown are as follows: HEAT Repeats (Red), FAT Domain (blue), RFB Domain (white), Kinase Domain (gray), FATC Domain (black). (c) Measurement in reduction of radial growth of *F. oxysporum* 6 strains to 50 ng/mL Rapamycin on minimal media plates amended with antibiotic compared to no-antibiotic MM plate controls. Three biological replicates were done for each treatment. Letters (a-d) indicate groups which are statistically different from one another using a T-test comparison between all groups. (d) Pictures of fungal growth on minimal media with and without antibiotic at 7 days post inoculation. Plates for the most resistant (Fo4287) and least resistant (Fo47) strains are shown.

Six out of the nine TOR paralogs encode full length proteins containing all functional domains (Figure 2b). One of the paralogous copies in the melon pathogen lacked the kinase domain and two paralogous copies (FOXG_15946 and FOTG_17386) had a truncation at the N-terminus (part of the HEAT repeats) of the protein. With the highly conserved domain structure, we were able to identify a shared mutation, M2345 to lysine, among all TOR paralogs near the very end of the catalytic loop of the kinase domain, while all other sequence motifs essential for TOR function were conserved (Figure 2b). Based on the protein structure, this catalytic loop forms the back of the ATP binding pocket, an invariant site for all conserved TOR kinases (45).

López-Berges et al. (46) reported that the truncation of FOXG_15946 was the result of a transposon insertion, and very low level of expression of this truncated copy was detected in rich medium. Our RNAseq data on the Arabidopsis pathogen Fo5176 during the course of infection also support the expression of both the ortholog (FOXB_14471) and the expanded copy (FOXB_02991) of the TOR kinases and the expression of the orthologous copy is roughly 3 times higher than the paralogous copy (Ma unpublished data).

As the TOR kinase is the direct target of the antibiotic rapamycin, we tested the sensitivity of 6 Fo strains, 4 with one TOR copy, and two with 2 TOR copies, to rapamycin. Rapamycin (50 ng/mL) reduced the rate of growth of all strains (p < 0.005) (Figure 2c, d). For the four strains containing the single copy of the TOR kinase, reduction in growth rate ranged from between 64% (MN25) to 82% (Fo47). The two strains carrying an additional TOR kinase, Fo5176 and Fo4287, showed increased resistance to the rapamycin treatment, resulting in 50% and 42% reduced growth, respectively. These results are consistent with an additional TOR kinase providing enhanced resistance to the drug rapamycin.

Functional studies suggested that the TOR signaling pathway may control pathogenic phenotypes, such as virulence in *F. oxysporum* infecting tomato (46) and toxin production in *Fusarium fujikuori* (47). Even though we don’t have direct evidence on the specific function of the expansion of TOR kinase among FOSC the high frequency (70% plant pathogenic strains), the preserved protein domain organization, and the increase resistance to rapamycin all suggest its potential functional involvement in the adaptation to diverse environments.

#### 3.2. Expansion of other signaling components involved in cell cycle control

TOR and its complex modulates cell growth patterns by partnering with other signaling components to control a number of regulatory subnetworks (48). Interestingly, we observed the expansion of several kinase families functioning as TOR partners, including CDC2, CLK and the BCK1 MAPK.

**BCK1:** The *Fusarium* BCK1 kinase family is expanded among 8 out of the 10 plant pathogenic isolates. All BCK1 kinases within FOSC fall into two clades, including an orthologous clade conserved in all *Fusarium* genomes, and a LS group only found within nine FOSC isolates (Supp. Fig. 4a). BCK1 is a member of the STE group kinase and is the only expanded family belonging highly conserved **category I** families (Figure 1d). Part of the MAPK signaling cascades under the control of the Rho1 GTPase and PKC1, BCK1 interacts with the TOR complex through direct binding with one of the subunits, LST8 (18, 20) and regulates transcription during the G1 to S phase transition as reported in *S. cerevisiae* (49).

**CMGC families:** Several kinase families of the group CMGC (Figure 1b), including CDC2, CDC2-like kinase (CLK), and SRPKL kinases, are expanded in FOSC genomes. Similar to TOR and BCK1 kinases, CLK gene expansion was only observed among FOSC genomes. An LS CLK was present in 7 out of 10 sequenced FOSC plant pathogenic strains. The core and the LS CLKs are phylogenetically distinct with strong bootstrap support (Sup. Fig. 4b). However, the expansion of CDC2 (Supp. Fig. 5a) and SRPKL kinases (Supp. Table 2) already occurred within the genu *Fusarium* before the split of *Fusarium* species. Collectively, SRPKL kinases constitute 4% to 7% of the total kinome among FOSC genomes (Supp. Table 2) and expansion happened multiple times. All other fungal genomes examined here contain a single CDC2 kinase, while *F graminearum, F. verticillioides* and all FOSC genomes have two. Expansion continued furthe within the FOSC. The tomato pathogen Fo4287 contains 10 additional LS CDC2 kinases and a pea pathogen FoHDV247 contains one extra copy. Not surprisingly, the two CDC2 gene detected in all *Fusarium* species are located in the core of the genome and recent duplicatioi events resulted in the 10 LS copies in the Fo4287 (2).

These expanded CMGC kinase families all have functions related to cell cycle contro directly or indirectly linked through the TOR signaling pathways. The CDC2 kinase is a majo regulator of the cell cycle. In *S. pombe*, the CDC2 kinase (SPBC11B10.09) directly regulates the G1-S to G2-M transitions and DNA damage repair (50, 51). CLK kinases are involved in variou cellular functions, including regulating the cell cycle in *S. pombe* (52), controlling ribosome an¢ tRNA synthesis in response to nutrient limitation and other cellular stresses in *S. cerevisiae* (53) and regulating cell wall biogenesis, vegetative growth, and sexual and asexual development in *Aspergillus nidulans* (54, 55). Like the CLK kinases, the SRPK kinases have been implicated in the control of SR protein mediated splicing in a TOR dependent manner in *S. cerevisiae* (56).

Functional importance of these expanded kinases in *Fusarium* genomes has been indicated by a few, however solid, functional studies. Deletion of the orthologous CDC2 kinases (FGSG_08468 in *F. graminearum* resulted in profound pleiotropic effects including reduced virulence an¢ decreased ascospore production. Deletion of the *Fusarium* specific copy (FGSG_03132) resulted in a milder, but still significant phenotype (15). In *F. graminearum*, the CLK kinase wa downregulated during conidial germination (15). Removal of one SRPKL kinase in *F. graminearum* (FGSG_02488 a SRPKL2) reduced DON production by half (15). The expression o the same gene was suppressed significantly during sexual development. Among the 9 core SRPKL kinases, 5 are expressed and four (two SRPKL1 kinases: FOXG_08977 and FOXG_10022 one SRPKL1 kinase: FOXG_21922; and one SRPKL3 kinase: FOXG_19803), were upregulated at 37°C. Two CDC2 kinases within the core were expressed at both conditions, while the expression of the 10 LS copies was not detected. Of the 51 Unclassified kinases in Fo4287, 12 core kinases and one LS kinase (FOXG_16175) had detectable expression in the RNAseq data generated from the reference strain.

#### 3.3. Enhanced environmental sensing accomplished through HisKs

One of the most significant kinase expansions occurs in the **HisK group** (Figure 3) known to play important roles in sensing and transducing many intra– and extracellular signals (36–39). The two yeast genomes, *S. cerevisae* and *S. pombe* encode one and three HisKs respectively, while the Hisk group consistently expands across the filamentous fungi (Figure 1d). All filamentous ascomycete fungal genomes included in this study have more than 10 HisKs and FOSC isolates have by far the most HisKs, with the tomato wilt pathogen Fol4287 and the tomato root rot pathogen FoHDV247 both predicted to have 23 HisKs. Based on a classical classification, fungal HisK are divided into 11 classes (36). Excluding *F. graminearum* which lacks the class IV HisK (TcsA kinase), all *Fusarium* genomes contain 10 distinct classes of HisK, only lacking the class VII that was reported in the *C. heterostrophus* and *Botrytis cinerea* genomes (36). Interestingly, the class II HisK is uniquely present in all *Fusarium* genomes and few other plant pathogenic Ascomycetes including *Cochliobolus heterostrophus* and *Bipolaris maydis* (36). The most significantly expanded FOSC HisKs are in class I and class IV (Figure 3), for instance class I HisKs continue to expand from 5 in *F. verticillioides* and 6 in *F. graminearum* to 7 or more among FOSC genomes.

**Figure 3:**
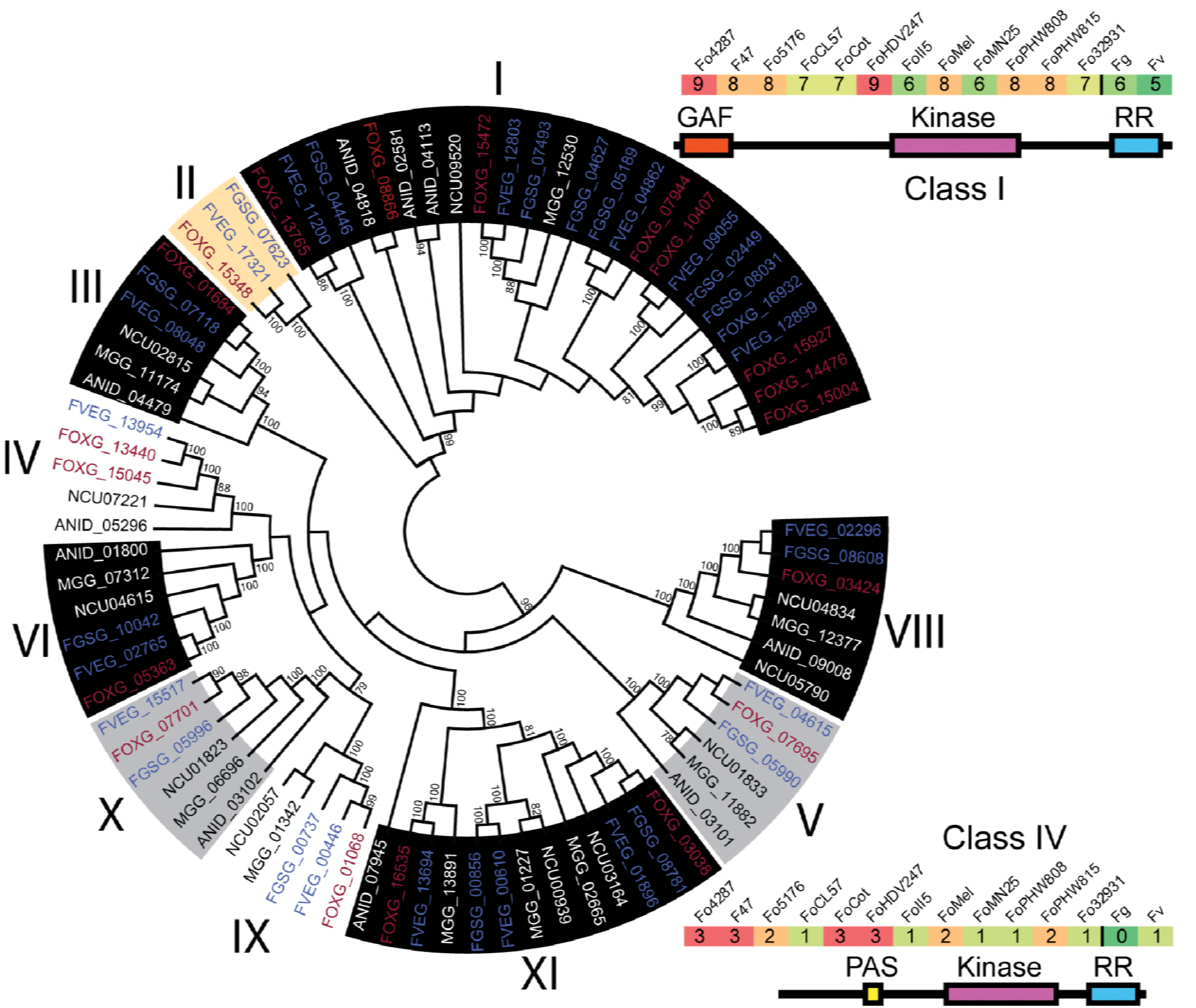
Histidine kinases are expanded in the Fusaria. A phylogenetic tree was constructed for all two-component signaling histidine kinase nucleotide sequences and labeled according to their class (family)(Neighbor Joining). Some kinases were excluded as they lacked a large portion of the aligned conserved sequence among HisK. Gray highlight, part of the conserved ascomycete kinome. Black highlight, part of the filamentous kinome. White highlight, part of the Fusaria kinome. Orange highlight, only found in Fusaria in this study. Fo gene IDs are colored red, Fg and Fv gene IDs are colored blue. All bootstrap values above 80% are shown. The domain structure for the class I and class IV HisK are shown along with a heatmap of the copy number for each class among the Fusaria.

According to protein domain organization, class I and class IV contain a GAF and a PAS N-terminal domain, respectively. PAS domains detect signals including light and oxygen, usually through binding to an associated cofactor (57). GAF domains form small binding pockets with potentials to bind to small signal molecules, such as cyclic GMP, cyclic AMP, or both. The motif is ubiquitously present in hundreds of signaling and sensory proteins from all kingdoms of life (58). Between class I HisKs, AA conservation of the GAF domain is low, likely indicating non-redundant function. Unlike the class I HisK, class IV copies have very similar N terminal domain sequences. Other than class I HisKs, class II, VIII, X and XI Hisks also contain GAF-related domains, while class IV, V, IX and XI also contain PAS-related domains.

First reported in prokaryotes, bacterial HisKs involve two-components where the first component senses environmental signals and autophosphorylates a conserved histidine residue, then transducing the signal to the second component through phosphate transfer (59). Among almost all eukaryotic systems, these two components are fused to form hybrid HisKs, containing the conserved histidine residue, ATP-binding and Response Regulator (RR) Receiver domains functioning in three-step “phosphorelay” reactions. Upon activation, the HisK phosphorylates itself twice, ending with phosphorylation of the response receiver domain at the C terminus. This phosphate is then transferred via a histidine phosphotransfer (Hpt) protein to a response regulator, which carries out downstream signaling functions. Each *Fusarium* isolate was found to have two RR proteins, orthologous to the yeast SKN7 and SSK1 RR which are both involved in the osmotic stress response downstream from the SLN1 HisK and YPD1 Hpt proteins (60, 61). The single yeast Hpt protein YPD1 has a single counterpart in *F. graminearum, F. verticillioides*, and each FOSC strain except for strains FoPHW808, FoPHW815, FoMelon, and FoCL57, which each have two. In these four cases, strains contain a copy which is located within the core genome and one which is located within the LS genome. Core genome copies share a 100% AA identity across the 143 AA protein, while LS genome copies share roughly 91% AA identity across 154 AA.

#### 3.4. Other expanded kinases

**HAL kinases:** Regulators of the cell’s primary potassium pumps (62, 63), the HAL kinases, are expanded in 6 out of the 12 sequenced FOSC genomes. In addition to the orthologous copies located in core genome regions, the second phylogenetically supported group contains all LS HAL kinases. Different from other FOSC specific expansions, this subfamily is also expanded in *M. oryzae* and *A. nidulans* (Supp. Fig. 5b). Noticeably, the HAL kinases are most significantly expanded to 6 copies in the strain Fo32931, a *F. oxysporum* strain isolated from an immunocompromised patient. Among the five HAL kinases in the reference genome Fo4287, all except one (FOXG_04685) were expressed (no change of expression with the shift of temperature), but the single core genome HAL kinase was expressed at least 30 fold higher than the LS HAL kinases.

**Unclassified protein kinases:** In addition, we observed a large number of kinases in the Unclassified family among all *Fusarium* genomes. On average, each *Fusarium* genome contains 41 unclassified kinases, accounting for roughly half of the kinome beyond the core ascomycete kinome and constituting 30 to 50 percent of LS kinome among FOSC genomes. No Unclassified kinases were present among all ascomycete fungal genomes examined, but four are conserved within the genus *Fusarium* (not shown) and more conservation was observed among FOSC genomes (Supp. Fig. 6). Among these unclassified kinases, eight were included in a *F. graminearum* knockout study and six mutants exhibit measurable phenotypes (15), including decreased the production of mycotoxin Deoxynivalenol in five mutants (FGSG_13509, FGSG_02153, FGSG_00792, FGSG_06420, FGSG_10591), and significant upregulation during sexual development for another (FGSG_12132).

**Atypical/FunK1:** Among the expanded families one of the least understood is the Atypical/Funk1, only found in complex Basidiomycota and Pezizomycota and possibly involved in the switch to multicellularity (64). Among Fusaria, copy number ranged from a single copy to 9, including one that is highly conserved among all *Fusarium* genomes. The expression of this conserved Funk1 gene in *F. graminearum* (FGSG_03499) is strongly upregulated during sexual development (15). Since no sexual reproduction was reported in *F. oxysporum*, the conservation of this gene within the *Fusarium* genus suggests functions other than sexual reproduction.

## Discussion

Kinases play key roles in transmitting external and internal signals and regulating complex cellular signaling responses. Genomes within the genus *Fusarium* have large kinomes compared to most fungi. As it is crucial for a pathogen to adapt to stresses encountered both outside and inside its host it is not surprising to see expansion of kinases among FOSC, as the species complex thrives in diverse hosts. The positive correlation between the total number of proteins and the size of the fungal kinome reported here was also reported in pathogenic *Microsporidia* species and several model organisms (10). Distinctively, this study defined a core kinome of the ascomycete fungi consisting of about 100 kinases, in agreement with a prediction based on gene to kinase count. This core kinome outlines the most fundamental kinase signaling pathways supporting the largest phylum of fungi.

Within the ascomycete fungal lineages some kinase families are more recalcitrant to change. The MAP Kinase cascades, Cell Kinase I, and Calcium/Calmodulin regulated kinases changed very little within the phylum over millions of years of evolutionary time. However, we observed the expansion of Cyclin-dependent Kinases and other close relatives, Histidine kinases, and Atypical kinases, suggesting their potential roles in species-specific adaptation along the various evolutionary trajectories of ascomycete fungi.

The number of Unclassified kinases increased significantly within the genus *Fusarium* and made up a majority of the kinases found within FOSC kinomes beyond those conserved amongst the filamentous fungi. In contrast, most non-Fusaria had relatively few Unclassified kinases. Additional research is necessary to understand how these Unclassified kinases contribute to *Fusarium* specific evolution and adaptation. However, it is clear that many Unclassified kinases are important, judging by the results from a reverse genetic screen (Wang et al. 2011), observed expression in our RNAseq data, and their conservation within the genus and across the FOSC.

Both the histidine kinase groups and the SPRKL family kinases were largely absent from the yeasts, moderately expanded in the *non-Fusarium* filamentous fungi, and appeared in large numbers in the FOSC. Little is known about the function of SRPKL kinases in fungal biology. HisKs are extensively used by bacteria and archaea to sense and respond to a variety of biotic and abiotic stimuli (65). In addition to regulating stress responses as reported in yeasts, the filamentous class III HisK were reported to regulate fungal morphogenesis and virulence in various human, plant, and insect pathogenic fungi (detailed review see: (37)). Overall, the function of histidine kinases is understudied in filamentous fungi. As HisKs are absent in mammals and some are essential for virulence in fungal pathogens, they represent interesting fungal targets for the discovery of new antifungal drugs. Although limited functional studies exist for Hisk or SRPKL kinases, the continuous expansion suggests functional importance of these understudied protein kinases. A better understanding of their functions would not only inform *Fusarium* biology, but could be extrapolated to other filamentous fungi and complex basidiomycetes.

Although LS kinases differ from their core counterparts, catalytic domains are generally conserved. For BCK1 kinases, the catalytic kinase domain of all nine LS BCK1 kinases is highly conserved (˜91% between groups), but all of them are roughly 800 AA shorter than their core genome counterparts, missing a portion of the N terminus. Similarly, sequences of the additional copies of CLK differ significantly from that of the orthologous copy, however the invariant residues within the catalytic kinase domain were conserved among all. In the case of CDC2 kinases, two of the three phosphorylation sites, Y15 and T14, are mutated in all LS CDC2 kinases. Since phosphorylation of these residues inhibits CDC2 kinase function (66), mutation at these two residues may lead to CDC2 kinase activity without tight control.

Most interestingly, we observed the repeated expansion of TOR kinase among the FOSC genomes. Most fungi have a single TOR protein, however two TOR paralogs were observed in both yeast genomes *S. cerevisiae* and *S. pombe* (43). Duplication of TOR kinases was also reported in *Batrachochytrium dendrobatidis*, an amphibian pathogenic chytrid (44). Seated at the center of many signal transduction pathways, TOR integrates the input from upstream pathways, sensing cellular nutrient, oxygen, and energy levels and dictates cellular responses. The convergent evolution toward TOR kinase duplication in the fungal kingdom might reflect selection for more finely tuned environmental response pathways. Interestingly we found that two FOSC strains containing an additional TOR copy were more resistant to rapamycin that those containing only one copy. Although our data for Fo4287 indicated that the TOR paralog was unexpressed, further studies will need to confirm this.

The expansion of families like the TOR kinase has been, in many cases, facilitated in part by the LS genome, that contributes to the unique and specific expansion of subfamilies, such as HisK, CLK, Atypical/FunK1, HAL, CMGC/SRPKL and Atypical/FRAP. Many of these kinases have either a known, or proposed, function in responding to environmental signals or cell cycle control and in many cases function downstream of TOR. Most expanded subfamilies can be linked to environmental signaling or cell cycle control; many under control of the TOR nutrient sensing complexes. Through the lenses of kinases, these pathogens seem to be enhancing their ability to sense their environment and tighten their regulation of the cell cycle, mediated primarily through the TOR signaling pathways.

## METHODS

### Generation of fungal kinomes

The two yeast genomes *Saccharomyces cerevisiae* S288C (Sc), *Schizosaccharomyces pombe* strain 972h- (Spom) were downloaded from NCBI. All other fungal genomes were downloaded from The Broad institute of MIT and Harvard and used for kinome analysis. Using the established Kinannote pipeline we generated the kinomes of the above fungal genomes. Briefly, Kinannote uses hidden Markov models generated from the manually aligned complete kinome of the slime mold *Dictyostelium discoideum* to search the given genome for kinases. It then identifies both well conserved eukaryotic protein kinases as well as unusual protein kinases. Finally Kinannote uses BLAST search results to classify the kinases based on family.

Kinannote assigned all kinases in all genomes into 135 different classes (Sup. Table 1). To create the conserved kinomes we removed any kinase subfamilies for which, under a given phylogenetic grouping, more than a single species was missing the family entirely. The groupings consisted of all species (Ascomycetes), all species except S. cerevisiae and S. pombe (Filamentous Fungi), only the genus *Fusarium* (Fusarium), and only members of the FOSC (FOSC). The number of conserved subfamily members was set as the lowest number among all species, excluding one.

### Gene alignment and BLAST

Fungal kinase protein and nucleotide sequences were downloaded from either The Broad institute or from the National Center for Biotechnology Information (NCBI). Sequences were aligned using either Muscle or Clustal through MEGA6 (67). Alignments were inspected manually and adjusted based on known conserved sequences. Sequence phylogeny was constructed using maximum likelihood and bootstrapped using 100 replicates.

BLAST was used to search for orthologs among the FOSC strains, conserved LS kinases, and to identify conserved regions of the LS genome. Homologs were considered top BLAST hits that had more than 90% nucleotide identity and covered the entire genomic gene sequence. To find groups of conserved LS kinases a file containing all LS kinase sequences from all strains was compared to an identical file using BLAST. A custom Perl script was generated to find and parse through results to identify groups of kinases which had significant BLAST results to each other. Identification of conserved LS regions was done by comparing each FOSC genome to the full LS gnome of Fo4287 including one core supercontig as a control for mapping (supercontig 14). BLAST results were filtered for regions matching greater than 90% nucleotide identify and alignment lengths greater than 3 kb. BLAST results were mapped to the Fo4287 LS reference sequence using BRIGS (68). Total overlap of each strains LS region with Fo4287 was calculated by summing all regions returned by BLAST.

### Generation of RNA samples and data analysis

*Fusarium oxysporum* f. sp. *lycopersici* 4287 gene expression during temperature stress was assayed using RNA-sequencing. *Fusarium oxysporum* spores (1×10^9^ spores) were cultured for 14 hours at 28°C (in the case of 28° growth experiment) or for 10 hours at 28°C and switched to 37°C for 4 hours (in the case of 37° growth experiment) in 200 ml of Minimal Medium supplemented with GluNa 25 mM and buffered with HEPES (20 mM final concentration) to pH 7.4 at 170 rpm. Three replicates of each condition were done for each species, with 36 RNA samples in total. Fungal tissue was collected with filter paper and RNA was extracted using a standard TRIzol RNA Isolation Reagent extraction (Life Technologies, Carlsbad CA). RNA Library was constructed using Illumina TruSeq Stranded mRNA Library Prep Kit (Illumina, CA) following manufacturer’s protocol and sequenced using the Illumina HiSeq platform (Illumina, CA).

Resulting data files were trimmed using Trimmomatic to remove poor quality reads (69). Reads were then aligned into BAM files using Rsubread (70). Gene expression and differentially expressed gene calls were then made using limma and edgeR (71, 72). Genes were called as expressed if both replicates had an RPKM value greater than 1.0. Genes were called as differentially expressed genes (DEGs) if the adjusted Pearson correlation value was less than 0.05.

### Rapamycin resistance screen

Cultures of Fo4287, Fo5276, Fo47, Foii5, and Fo32931 were grown in Potato Dextrose Broth (Becton, Dickinson and Company, Sparks, MD), the strain MN25 was grown on Potato Dextrose Agar (Becton, Dickinson and Company, Sparks, MD), for 5 days and spotted into the center of either minimal media plates (73) or minimal media plated with a final concentration of 50 ng/mL rapamycin. Plates were stored at 28 degrees for 48 hours and then transferred to room temperature. After transfer to room temperature the diameter of each colony was measured at 24 hour intervals. Growth rate was determined as the average increase in size, in millimeters, per 24 hours. Reduction in growth rate was calculated as (100 – ((growth rate on rap. plate)/(growth rate on control plate)*100)). In order to generate a range of error, all 9 comparisons of growth rate between the 3 control plates and 3 rapamycin plates were used to calculate reduction in growth. All species/condition plates were done in triplicate.

## Data Availability

The *Fusarium oxysporum* temperature RNAseq data for this study has been deposited into the NCBI GEO repository under the accession number GSE113332.

## Author contributions

Project Design, data generation and data analysis: GAD, JG, YZ, HCK and LJM.

Manuscript writing: GAD and LJM.

RNA sample prep and data analysis: LG and GAD.

## Acknowledgement

This project was supported by the National Research Initiative Competitive Grants Program Grant no. 2008-35604-18800 and MASR-2009-04374 from the USDA National Institute of Food and Agriculture. Data analysis was conducted at the MGHPCC. The funding for the RNAseq was provided by the National Research Initiative Hatch Grants Program Grant no. MAS00441. LJM is also supported by Investigator Award in Infectious Diseases and Pathogenesis by the Burroughs Wellcome Fund BWF-1014893. GAD and LJM are also supported by the National Science Foundation ISO-165241.

---

**Supplementary Figure 1: The LS genome contributes to kinome expansion.**(a) The number of genes in the LS genome was plotted against the number of kinases found with the LS genome of each Fo strain. (b) The LS genome of each FOSC strain was compared to the LS genome of the reference strain. Colored lines indicate regions of conservation. Genomes were mapped to core genome contig 14 (roughly the first 1/5^th^ of chromosome 1, a conserved chromosome) as a control. Large regions of color indicate potentially shared LS genome contigs. Names listed at the top of the ring correspond to the strain Fo strain for that particular colored ring (i.e. FoCotton is listed as Cotton). For brevity the Fo identifier was left off of the strain ID labels.

**Supplementary Figure 2: Nucleotide conservation among TOR kinases confirms single origin for LS TOR kinases.**(a) The domain structure of the TOR kinase is shown with the M2345L marked as a red band. Above the graph indicates the percent amino acid conservation in 10 bp windows across a gap deleted protein sequences for Fg, Fv, and Fo4287 (purple), all FOSC core genome TOR kinases (blue), and FOSC LS genome TOR kinases (black). (b) The nucleotide identity between TOR kinases in Fusaria was compared. Gene IDs are organized into LS, Core, and Fg and Fv at the bottom. Colors are relative representations of the nucleotide identity.

**Supplementary Figure 3: Differential inhibition of FOSC isolate growth by Rapamycin.**(top) Images for one representative pair of plates were taken for each strain at 7 dpi. Strains were cultured on minimal media (top plate) or minimal media with 50 ng/mL Rapamycin (bottom plate). (below) The growth rate in millimeters per 24 hours was calculated for each strain on both media. All MM and MM+Rap pair differences are statistically significant (t-test, P.value < 0.005). All plates were done in triplicate.

**Supplementary Figure 4: BCK1 and CLK Kinase Sub-families.**Phylogenetic trees (Maximum Likelihood) using nucleotide sequences from BCK1 (a) and CLK (b) kinase sub-families from all species show the clear separation of core and LS genome orthologs. Scale bar is substitution rate. Black boxes, LS genome genes. White boxes, core genome genes. Question marks, unknown contigs.

**Supplementary Figure 5: CDC2 and HAL Kinase Sub-families.**Phylogenetic trees (Maximum Likelihood) using nucleotide sequences from CDC2 (a) and HAL (b) kinase sub-families from all species show the clear separation of core and LS genome orthologs. Scale bar is substitution rate. Black boxes, LS genome genes. White boxes, core genome genes. Question marks, unknown contigs.

**Supplementary Figure 6: Shared Unclassified kinases among the FOSC.**Phylogenetic trees (Maximum Likelihood) using nucleotide sequences from all FOSC unclassified kinases. Bars around the outside of the tree denote clades supported with high bootstrap values. When 2 clades are adjacent one is colored blue to aid visually. The number next to each bar indicates the number of FOSC strains which contain a gene from that clade.

